# An innovative in vitro model for studying the biology of cardiac fibroblasts originating from the epicardium

**DOI:** 10.1101/2025.05.23.655755

**Authors:** Claudia Müller-Sánchez, María Gertrudis Muñiz-Banciella, Manuel Reina, Francesc X. Soriano, Ofelia M. Martínez-Estrada

## Abstract

Embryonic epicardium is a major source of cardiac fibroblasts (CFs), which play essential roles in heart development and response to heart injury. In this study, we developed a novel mouse model to identify distinct populations of epicardium-derived CFs. Our Wt1^GFP/+^;Wt1Cre;ROSA26-tdRFP model enables lineage tracing of WT1Cre-labeled (RFP^+^) fibroblasts and the identification of cells actively expressing WT1 (GFP^+^). Flow cytometry at early postnatal stages showed that RFP^+^ cells form a heterogeneous stromal population, with 20.13% co-expressing GFP, indicating persistent WT1 expression in a subset. We successfully immortalized RFP^+^ cardiac stromal cells, highly enriched in fibroblasts, by excluding other Wt1Cre-active cell types. Through culture condition optimization, we could selectively expand or differentiate specific fibroblast subpopulations, increasing the model’s utility. These immortalized cells, carrying an integrated WT1 reporter system, provide a robust in vitro platform to study fibroblast activation, differentiation, and plasticity under defined conditions.

**Summary blurb:** This study presents a novel in vitro mouse model for investigating the activation, identity, and functional properties of epicardium-derived cardiac fibroblasts.

## INTRODUCTION

Cardiac fibroblasts (CFs) are essential non-myocyte cells that play a crucial role in heart development and homeostasis (Tallquist, 2018). They are among the most abundant non-cardiomyocyte cell types in the postnatal heart and are responsible for extensive extracellular matrix (ECM) deposition. Beyond their structural role, CFs produce paracrine factors critical to heart development, myocardial growth, functional maturation, and blood vessel formation (Acharya et al, 2012; Hortells et al, 2020; Ieda et al, 2009). They are also central to the formation of pathological fibrosis, a hallmark of various cardiovascular diseases, by promoting excessive ECM deposition and tissue remodeling (Humeres & Frangogiannis, 2019).

The understanding of CF biology has advanced significantly over the past decade (Tallquist & Molkentin, 2017). Although several origins have been proposed, lineage tracing studies in mice have unequivocally demonstrated that adult resident CFs primarily arise from the embryonic epicardium and endocardium, with the epicardium being the predominant source (Moore-Morris et al, 2014; Tallquist, 2018). Notably, the population of resident CFs of epicardial origin expands in response to adult cardiac fibrosis, such as that induced by pressure overload, fibroelastosis, or myocardial infarction (Forte et al, 2020; Moore-Morris et al, 2014; Ruiz-Villalba et al, 2015; Zhang et al, 2017).

Compared with other cell types, the absence of unique fibroblast markers, coupled with their inherent heterogeneity, has made the isolation of fibroblasts particularly challenging when using conventional techniques such as fluorescence-activated cell sorting (FACS) (Tallquist & Molkentin, 2017). Various genetic mouse models have, therefore, been developed to facilitate fibroblast identification. Some models employ direct fibroblast reporter systems, such as the Pdgfra-GFP and Collagen1a1-GFP reporter lines (Hamilton et al, 2003; Yata et al, 2003). Additionally, Cre transgenic mouse models have significantly improved the identification and characterization of CFs (Acharya et al, 2011; Tallquist & Molkentin, 2017; Wessels et al, 2012; Zhang et al, 2008; Zhou et al, 2008).

The WT1 gene encodes a zinc finger protein essential for heart development (Hastie, 2017; Moore et al, 1999). WT1 is highly expressed in the epicardium and epicardial derived cells (EPDCs) during heart development, where it plays a critical role in their formation and function (Martinez-Estrada et al, 2010; Moore et al, 1999; Velecela et al, 2019). Epicardial derivatives, such as CFs, also express WT1 both during heart development and following myocardial infarction (Braitsch et al, 2013; Perez-Pomares et al, 2002; Zhou et al, 2011).

These data, originally identified by immunostaining, have been validated by several transcriptomic studies(Deng et al, 2025; Feng et al, 2022). Both constitutive and inducible Wt1Cre mouse models have been generated and used to study fibroblast formation and expansion during development (Wessels et al, 2012; Zhou et al, 2008), and these models have become essential tools for investigating resident fibroblasts of epicardial origin in heart diseases (Forte et al, 2020; Moore-Morris et al, 2018; Ruiz-Villalba et al, 2015).

In this study, we used a constitutive BAC Wt1Cre transgenic mouse model, which efficiently labels resident CFs, and generated a new Wt1^GFP/+^;Wt1Cre;ROSA26-tdRFP mouse model. This includes WT1 lineage cells that are RFP-positive (RFP^+^), while cells actively expressing WT1 are GFP-positive (GFP^+^). We subsequently established immortalized CFs from this model, which represent a valuable and versatile in vitro tool for dissecting the molecular mechanisms underlying fibroblast activation, differentiation, and plasticity.

## RESULTS

### Characterizing cardiac cells from the Wt1^GFP/+^;Wt1Cre;ROSA26-tdRFP mouse model

Although several studies have used Wt1Cre mouse models to study fibroblast formation during cardiac development and disease states, none have combined lineage tracing with a Wt1 reporter model to identify both Wt1Cre progeny and Wt1-expressing cells simultaneously, which can be used to FACS-isolate different populations of Wt1Cre positive cells (Forte et al, 2020; Moore-Morris et al, 2014; Ruiz-Villalba et al, 2015; Wessels et al, 2012). We aimed to generate such a model by crossbreeding Wt1Cre;ROSA26-tdRFP mice with Wt1^GFP/+^ mice (Fig. 1A).

**Figure 1.**
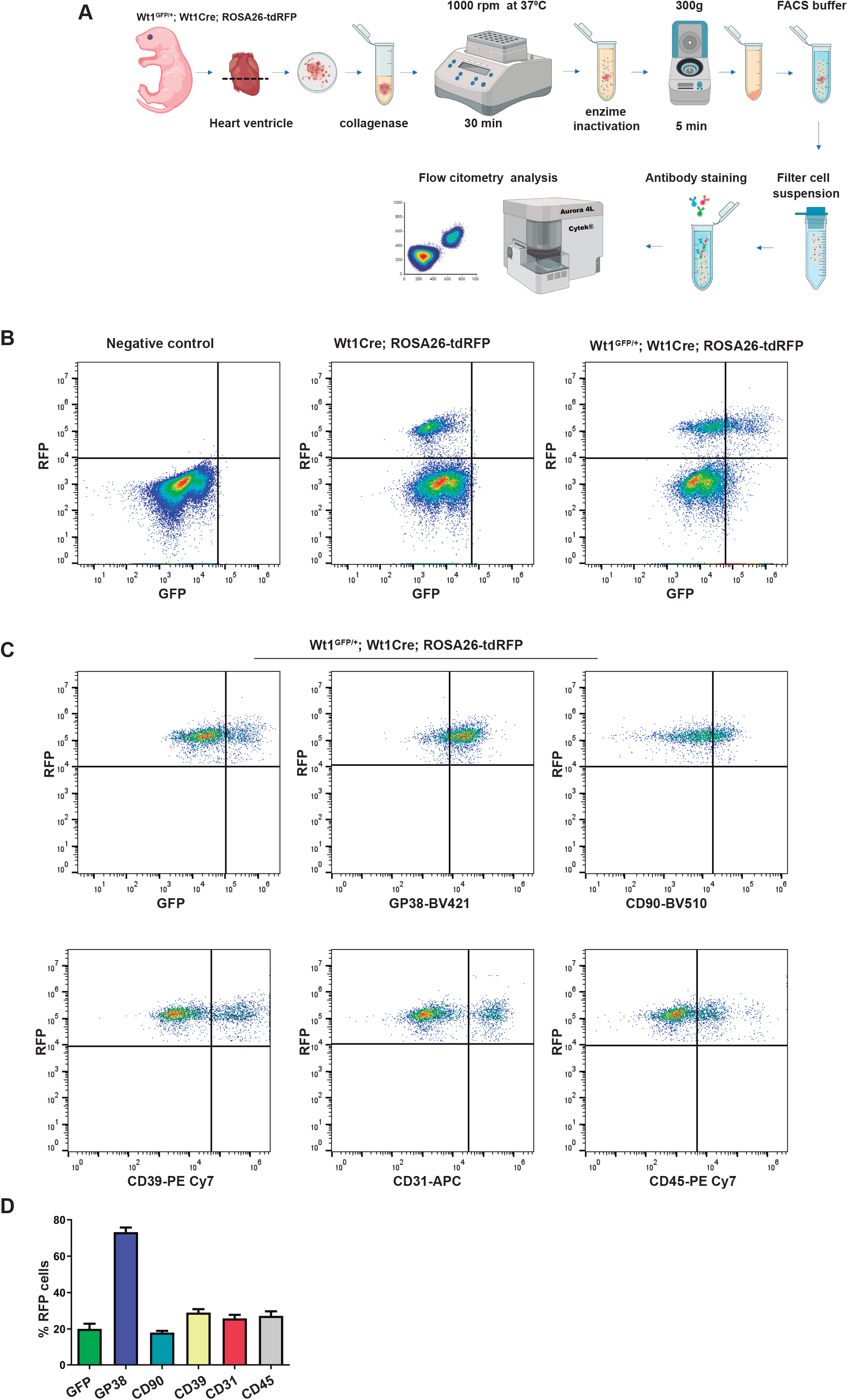
Generation and characterization of freshly isolated cardiac cells from the Wt1^GFP/+^;Wt1Cre;ROSA26-tdRFP mouse model. **(A)** Overview of the protocol for enzymatic digestion and flow cytometry analysis of heart ventricles from Wt1^GFP/+^;Wt1Cre;ROSA26-tdRFP. **(B)** Flow cytometry analysis of enzymatically digested P2 heart ventricles from Wt1^GFP/+^;Wt1Cre;ROSA26-tdRFP mice reveals that a subset of RFP^+^ cells retains sustained GFP expression. Littermate negative mice were used to establish gating parameters. **(C, D)** Representative flow cytometry plots for identification of RFP^+^ subsets and corresponding quantification. Data are presented as mean ± s.e.m. (n = 4).

In the Wt1^GFP/+^ model, GFP is knocked into exon 1 and driven by the endogenous Wt1 regulatory sequences, providing a GFP reporter that faithfully reflects Wt1 expression (Hosen et al, 2007; Velecela et al, 2019). In the new model, Wt1^GFP/+^;Wt1Cre;ROSA26-tdRFP mice contain RFP-positive (RFP^+^) cells that exhibiting Cre recombinase activity, along with their progeny, while GFP-positive (GFP^+^) cells express abundant WT1 levels. Flow cytometry analysis of ventricular cardiac cells at early postnatal stages showed that RFP^+^ cells represented 13.75% of the total cardiac cell population. Of these, 20.13% were GFP^+^, indicating sustained WT1 expression in a cell subset (Fig. 1A, B, Table 1,2).

**Table 1.**
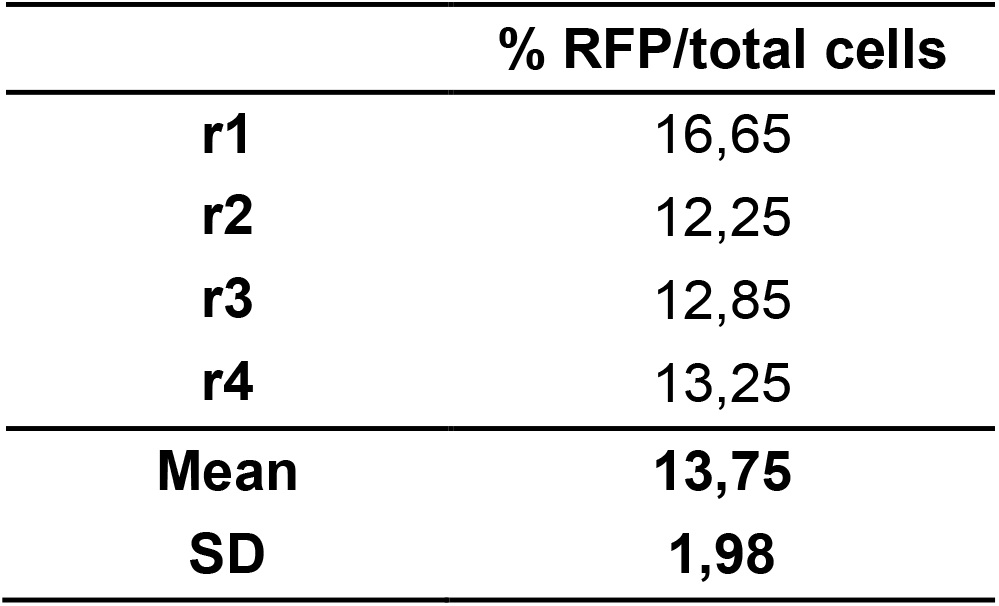
Percentage of RFP-positive cells relative to the total number of cells, determined by flow cytometry. Data are mean ± s.e.m. (n=4).

**Table 2.** Percentage of cells expressing the indicated markers within the RFP-positive cell population, determined by flow cytometry. Data are mean ± s.e.m. (n=4).

| RFP+ cells |  |  |  |  |  |  |
| --- | --- | --- | --- | --- | --- | --- |
|  | **GFP** | **GP38** | **CD90** | **CD39** | **CD31** | **CD45** |
| **r1** | 20,5 | 79,90 | 19,4 | 30,50 | 22,60 | 24,50 |
| **r2** | 24,1 | 69,50 | 19,9 | 30,90 | 24,70 | 24,50 |
| **r3** | 23,8 | 69,20 | 15,8 | 31,30 | 24,50 | 25,40 |
| **r4** | 12,10 | 74,40 | 17 | 23,40 | 31,50 | 34,60 |
| **Mean** | **20,13** | **73,25** | **18,03** | **29,03** | **25,83** | **27,25** |
| **SD** | **5,59** | **5,03** | **1,95** | **3,76** | **3,90** | **4,92** |

To further characterize the RFP^+^ population, we employed a panel of several antibodies. This showed that 25.83% of RFP^+^ cells expressed CD31 and 27.25% expressed CD45, confirming the presence of endothelial and hematopoietic cells within the RFP^+^ population, consistent with previous reports (Fig. 1C, D, Table 2) (Cano et al, 2016; Pogontke et al, 2022). Notably, 69.61% of the RFP^+^ CD45^+^ cells and 76.34% of the RFP^+^ CD31^+^ cells co-expressed GFP (Table 3). We also included markers commonly used to identify the stromal cell populations GP38, CD90, and CD39. Among the RFP^+^ cells, 73.25% were GP38^+^, 18.03% were CD90^+^, and 29.03% were CD39^+^ (Fig. 1C, D, Table 2). Of note, the percentages of GFP^+^ cells varied among these populations, accounting for 28.91% of GP38^+^, 36.46% of CD90^+^, and 71.04% of CD39^+^ cells (Table 4).

**Table 3.**
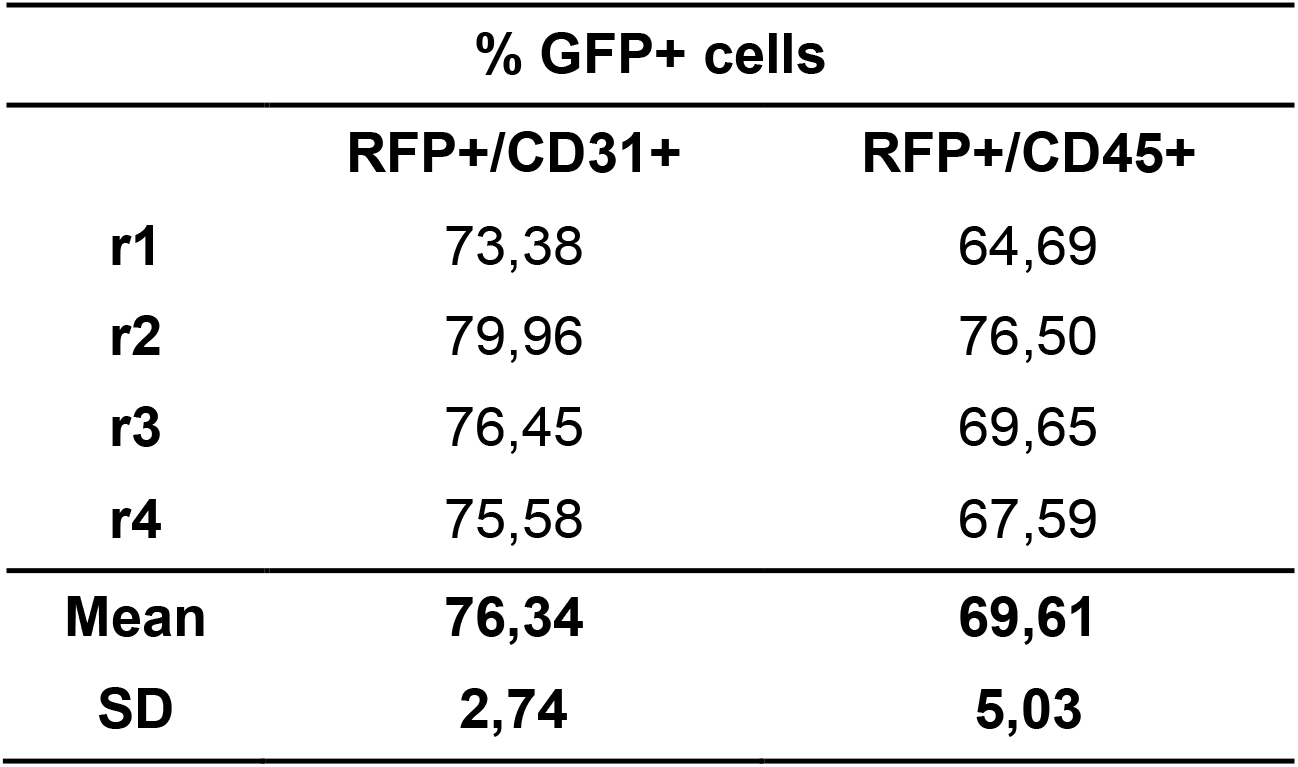
Percentage of GFP cells within the RFP+ CD31+ and RFP+ CD45+ cell population, determined by flow cytometry. Data are mean ± s.e.m. (n=4).

**Table 4.** Percentage of cells expressing the indicated markers within the RFP-positive cell population, determined by flow cytometry. Data are mean ± s.e.m. (n=4).

|  | RFP+/GP38+/- |  |  | RFP+/CD90+/- |  |  | RFP+/CD39+/- |  |  |
| --- | --- | --- | --- | --- | --- | --- | --- | --- | --- |
|  | GFP | CD90 | CD39 | GFP | GP38 | CD39 | GFP | GP38 | CD90 |
| **r1** | 29,69 | 22,92 | 27,87 | 38,94 | 85,86 | 34,43 | 70,97 | 77,76 | 21,82 |
| **r2** | 37,09 | 23,18 | 24,90 | 46,52 | 76,14 | 45,66 | 78,65 | 57,96 | 8,68 |
| **r3** | 30,81 | 18,69 | 24,54 | 32,06 | 75,82 | 28,65 | 77,70 | 58,93 | 9,44 |
| **r4** | 18,06 | 20,23 | 16,95 | 28,33 | 80,06 | 38,28 | 56,84 | 57,67 | 18,73 |
| **Mean** | **28,91** | **21,25** | **23,56** | **36,46** | **79,47** | **36,76** | **71,04** | **63,08** | **14,67** |
| **SD** | **7,93** | **2,17** | **4,65** | **8,02** | **4,68** | **7,13** | **10,07** | **9,80** | **6,61** |

Taken together, these results demonstrate that the Wt1^GFP/+^;Wt1Cre;ROSA26-tdRFP mouse model allows the identification of different populations of Wt1Cre-positive stromal cells based on GFP expression.

### Immortalization of cardiac cells in the Wt1^GFP/+^;Wt1Cre;ROSA26-tdRFP mouse model

Most studies using CF cultures rely on selective attachment to isolate the cell population, often without additional enrichment strategies (Garvin & Katwa, 2023). In this study, we FACS-isolated RFP^+^ ventricular cardiac stromal cells from the Wt1^GFP/+^;Wt1Cre;ROSA26-tdRFP mouse model, excluding endothelial (RFP^+^, CD31^+^, CD45^−^), hematopoietic (RFP^+^, CD31^−^, CD45^+^), and high GFP-expressing cells, thereby including epicardial cells, among others (Fig. 2A). The sorted RFP^+^ cells were cultured on poly-L-lysine-coated plates, where they exhibited a flattened, irregular morphology with abundant cytoplasmic protrusions (Fig. 2B). Next, the RFP^+^ cells were immortalized using a viral vector containing the SV40 large T antigen (SV40 TAg), which is well-recognized for its robust cell immortalization capability. Following puromycin selection, the immortalized cells exhibited a more uniform morphology with abundant cytoplasmic protrusions (Fig. 2C). Immunofluorescence analysis confirmed uniform expression of vimentin, a key fibroblast-specific marker, in all cells, along with the presence of some patches of smooth muscle actin (SMA)-positive cells (Fig. 2D, E).

**Figure 2.**
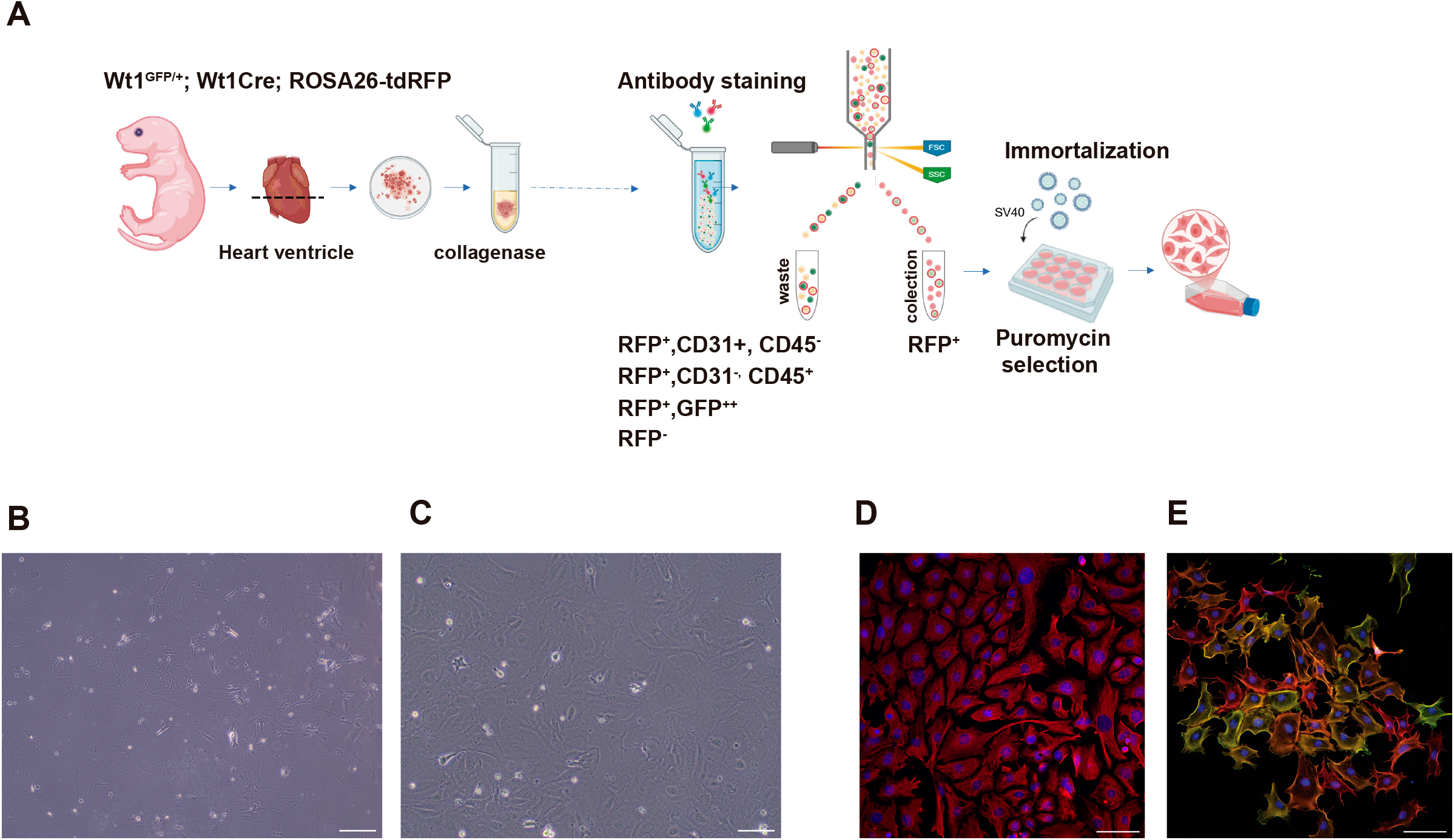
Generation of immortalized cardiac cells from Wt1^GFP/+^;Wt1Cre;ROSA26-tdRFP mice. **(A)** Workflow for the strategy used to generate and immortalize postnatal RFP^+^ cardiac fibroblasts from Wt1^GFP/+^;Wt1Cre;ROSA26-tdRFP mice excluding endothelial cells (RFP^+^,CD31^+^, CD45^-^), hematopoietic cells (RFP^+^CD31^-^,CD45^+^), and cells with the highest GFP expression (RFP^+^,GFP^++^). **(B, C)** Bright-field image of primary (B) and immortalized (C) cardiac fibroblasts. **(D)** Immunofluorescence staining for the intermediate filament vimentin, demonstrating abundant vimentin expression in immortalized cells. **(E)** Immunofluorescence staining for SMA (green) and phalloidin (red), exhibiting some patches of SMA-positive cells. Scale bars: 200 µm in (B, C); 50 µm in (D, E).

These results validate the successful immortalization of FACS-sorted RFP^+^ cardiac stromal cells.

### Characterization of cardiac RFP^+^ immortalized cells derived from the Wt1^GFP/+^;Wt1Cre;R26-tdRFP mouse model

Generating immortalized cells allows the study of CF signaling, proliferation, and activation with less need to use animals. This can be achieved because the immortalized cells remain proliferative across multiple passages without showing signs of senescence. Next, we aimed to characterize the RFP-immortalized cells in detail. Flow cytometry confirmed uniform RFP expression across all cells, with 15.49% expressing GFP (Fig. 3A, Table 5). When we included the antibodies used for in vivo analysis, the flow cytometry revealed that 97.40% of the RFP^+^ immortalized cells expressed GP38, 63.12% expressed CD90, and 5.87% expressed CD39 (Fig. 3A, Table 5). Consistent with in vivo observations, the proportion of GFP^+^ cells varied among subpopulations: 10.14%, 11.61%, and 29.98% among the RFP^+^GP38^+^, RFP^+^CD90^+^, and RFP^+^CD39^+^ subpopulations, respectively (Fig. 3B, Table 6). Given that none of these markers are specific to fibroblasts, and that previous studies have shown the Wt1Cre line can recombine in other cell types, we assessed the expression of genes enriched in other cardiac cell populations, including *Des* (smooth muscle cells), *Rgs5* (pericytes), and *Myh6* (cardiomyocytes) (Fig. 3C). We also examined the expression of genes highly expressed in the epicardium, such as *Msln* and *Upk3b* (Fig. 3D). The qRT-PCR analysis revealed that immortalized and sorted RFP^+^ cells exhibited undetectable levels of these non-fibroblast cell markers and abundant *Vim* expression (Fig. 3C, D). Since the vast majority of the immortalized cells were GP38^+^CD39^−^, we FACS-sorted four distinct cell subpopulations based on CD90 and GFP expression: CD90^+^GFP^+^, CD90^+^GFP^−^, CD90^−^GFP^+^, and CD90^−^GFP^−^. Next, we decided to compare the potential contribution of each of these cell populations to the SMA-positive patches observed in the cell culture. Interestingly, qRT-PCR analysis revealed that the CD90^+^GFP^−^ subpopulation expressed significantly higher levels of *Acta2* compared to the other subpopulations (Fig. 3E, F).

**Table 5.**
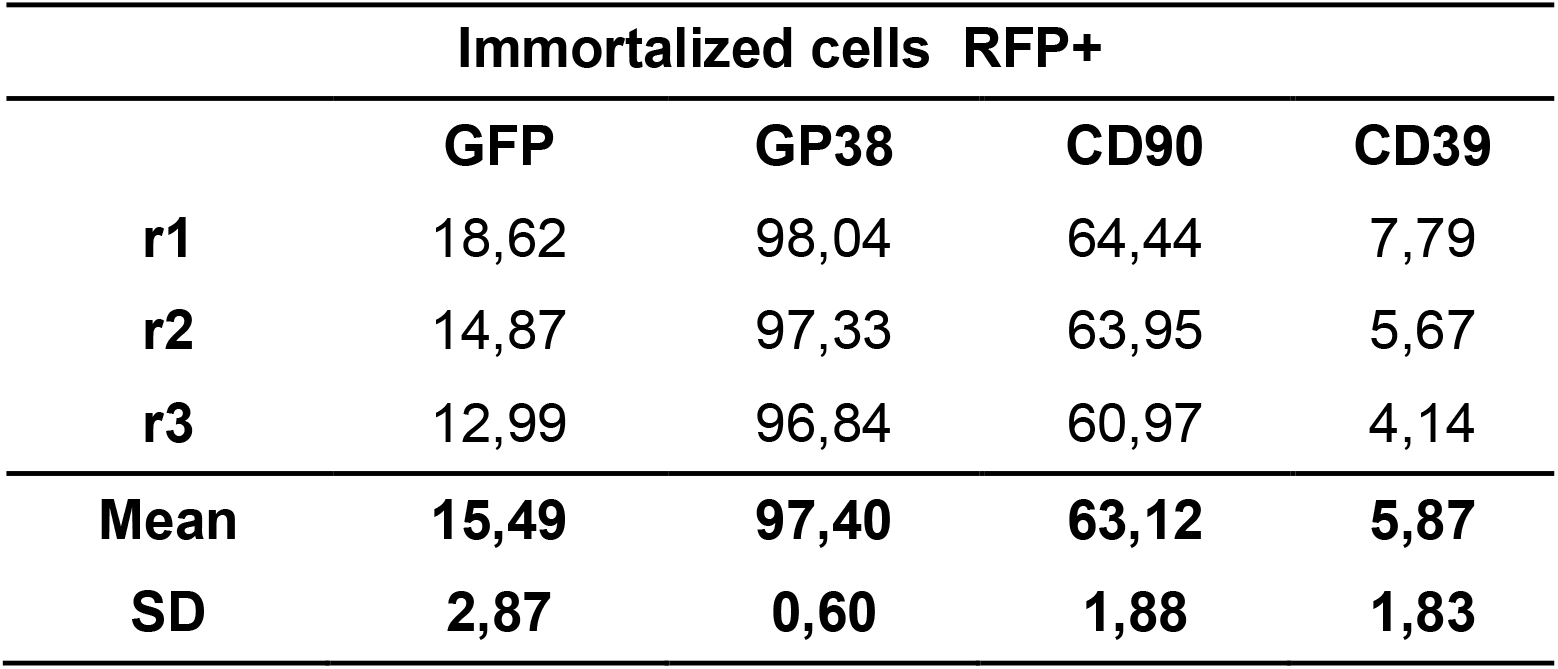
Percentage of immortalized cells expressing the indicated markers within the RFP-positive cell population, determined by flow cytometry. Data are mean ± s.e.m. (n=3).

**Table 6.**
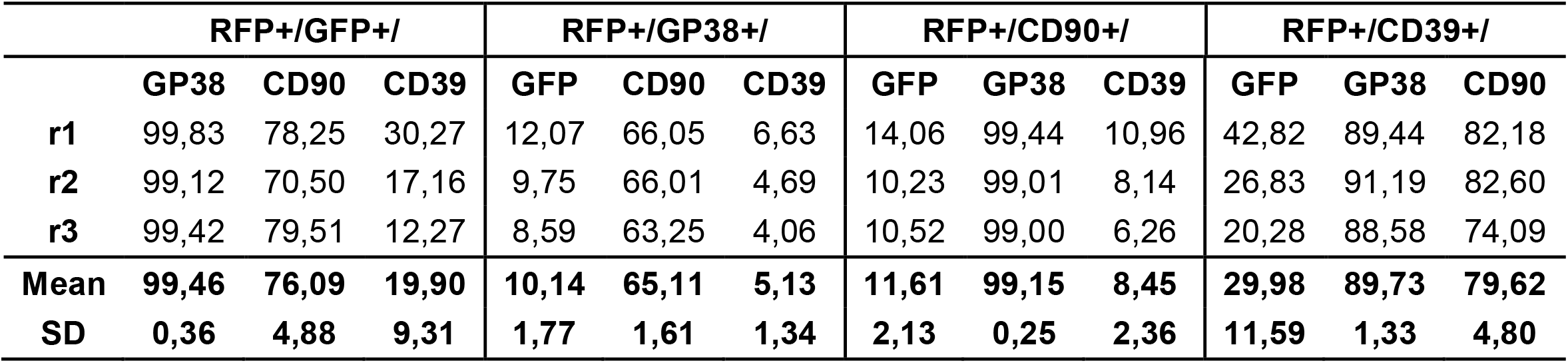
Percentage of immortalized cells at expressing the indicated markers within the RFP-positive cell population, determined by flow cytometry. Data are mean ± s.e.m. (n=3).

|  | RFP+/GFP+/- |  |  | RFP+/GP38+/- |  |  | RFP+/CD90+/- |  |  | RFP+/CD39+/- |  |  |
| --- | --- | --- | --- | --- | --- | --- | --- | --- | --- | --- | --- | --- |
|  | GP38 | CD90 | CD39 | GFP | CD90 | CD39 | GFP | GP38 | CD39 | GFP | GP38 | CD90 |
| **r1** | 99,83 | 78,25 | 30,27 | 12,07 | 66,05 | 6,63 | 14,06 | 99,44 | 10,96 | 42,82 | 89,44 | 82,18 |
| **r2** | 99,12 | 70,50 | 17,16 | 9,75 | 66,01 | 4,69 | 10,23 | 99,01 | 8,14 | 26,83 | 91,19 | 82,60 |
| **r3** | 99,42 | 79,51 | 12,27 | 8,59 | 63,25 | 4,06 | 10,52 | 99,00 | 6,26 | 20,28 | 88,58 | 74,09 |
| **Mean** | **99,46** | **76,09** | **19,90** | **10,14** | **65,11** | **5,13** | **11,61** | **99,15** | **8,45** | **29,98** | **89,73** | **79,62** |
| **SD** | **0,36** | **4,88** | **9,31** | **1,77** | **1,61** | **1,34** | **2,13** | **0,25** | **2,36** | **11,59** | **1,33** | **4,80** |

**Figure 3.**
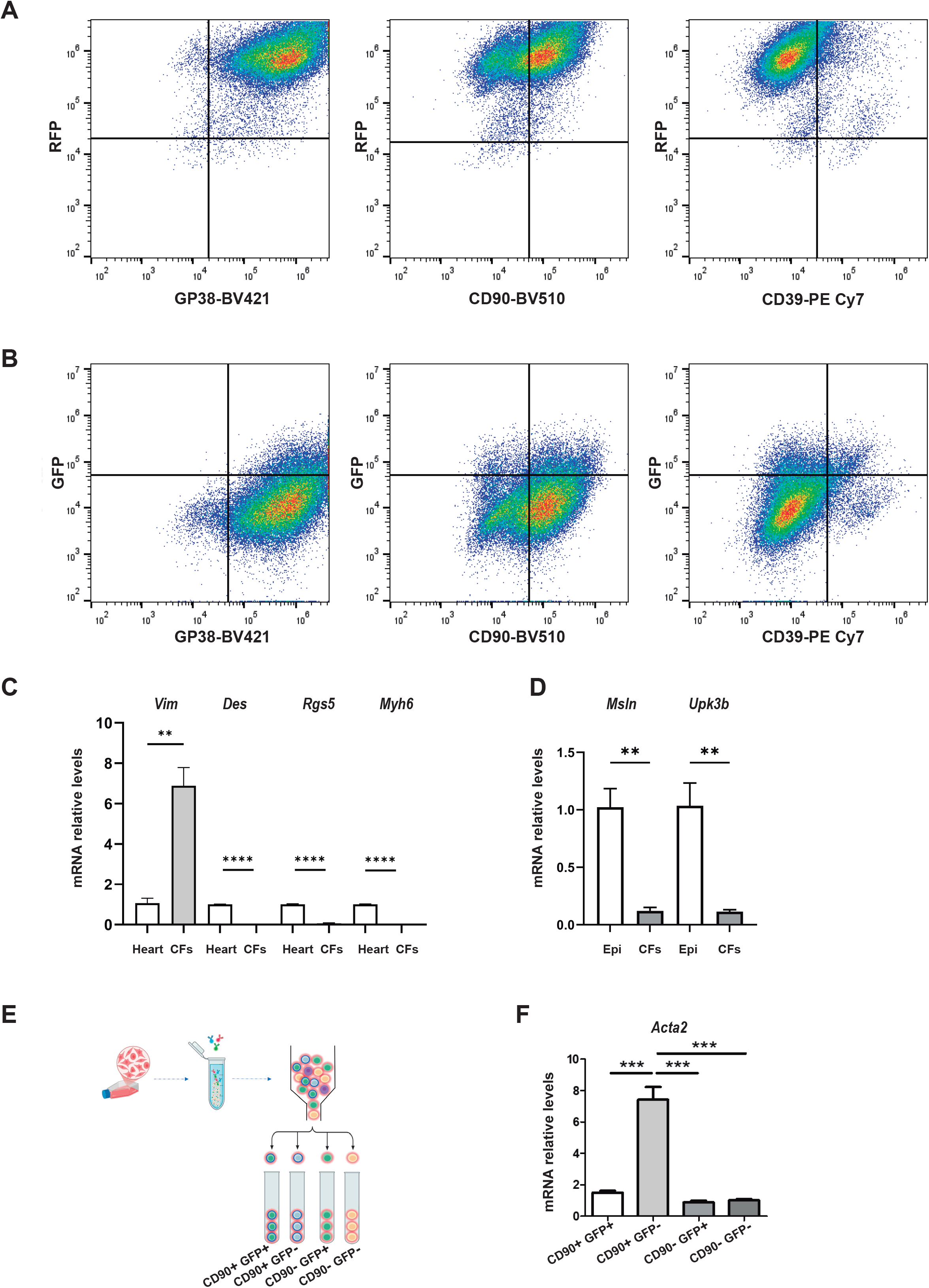
Characterization of immortalized cardiac fibroblasts from Wt1^GFP/+^;Wt1Cre;ROSA26-tdRFP mouse model. **(A)** Representative flow cytometry plots showing the identification of RFP^+^ cardiac fibroblast subsets **(B)** Representative flow cytometry plots for identification of GFP^+^ subsets. **(C, D)** qRT-PCR analysis of the indicated genes in immortalized cardiac cells from Wt1^GFP/+^;Wt1Cre;ROSA26-tdRFP mice. Compared with total heart tissue or immortalized epicardial cells (Epi), immortalized fibroblasts exhibited undetectable expression of markers specific to smooth muscle cells, pericytes, cardiomyocytes, and epicardial cells. Data are presented as mean ± s.e.m. (n = 3); **P < 0.001, ***P < 0.0001 two-tailed student t-test. **(E)** Schematic representation of the FACS-based isolation of four fibroblast subpopulations defined by CD90 and GFP expression. **(F)** qRT–PCR analysis of Acta2 expression in these subsets revealed elevated Acta2 levels in the CD90^+^GFP^−^ population. Data are presented as mean ± s.e.m. (n = 6); ***P < 0.0001, One-way ANOVA followed by Tukey’s post-hoc test.

Collectively, these findings demonstrate that the RFP^+^ cells represent a heterogeneous population of CFs.

### Transforming growth factor-beta (TGF-β) modulates the fibrotic response in RFP^+^ immortalized CFs

Increased SMA expression is a widely used marker for identifying fibroblast activation (Garvin & Katwa, 2023). After observing SMA-positive patches in our culture, we sought conditions to prevent this activation. Commercial fibroblast-specific media reduce serum concentration, which helps prevent spontaneous activation. As expected, CFs cultured in this medium exhibited lower *Acta2* expression compared to those cultured in DMEM (Fig. 4A). Interestingly, this reduction in *Acta2* expression was accompanied by an increase in the CD90^+^GFP^+^ and CD90^−^GFP^+^ populations, along with a decrease in the CD90^+^GFP^−^ and CD90^−^GFP^−^ populations (Fig. 4B, C).

**Figure 4.**
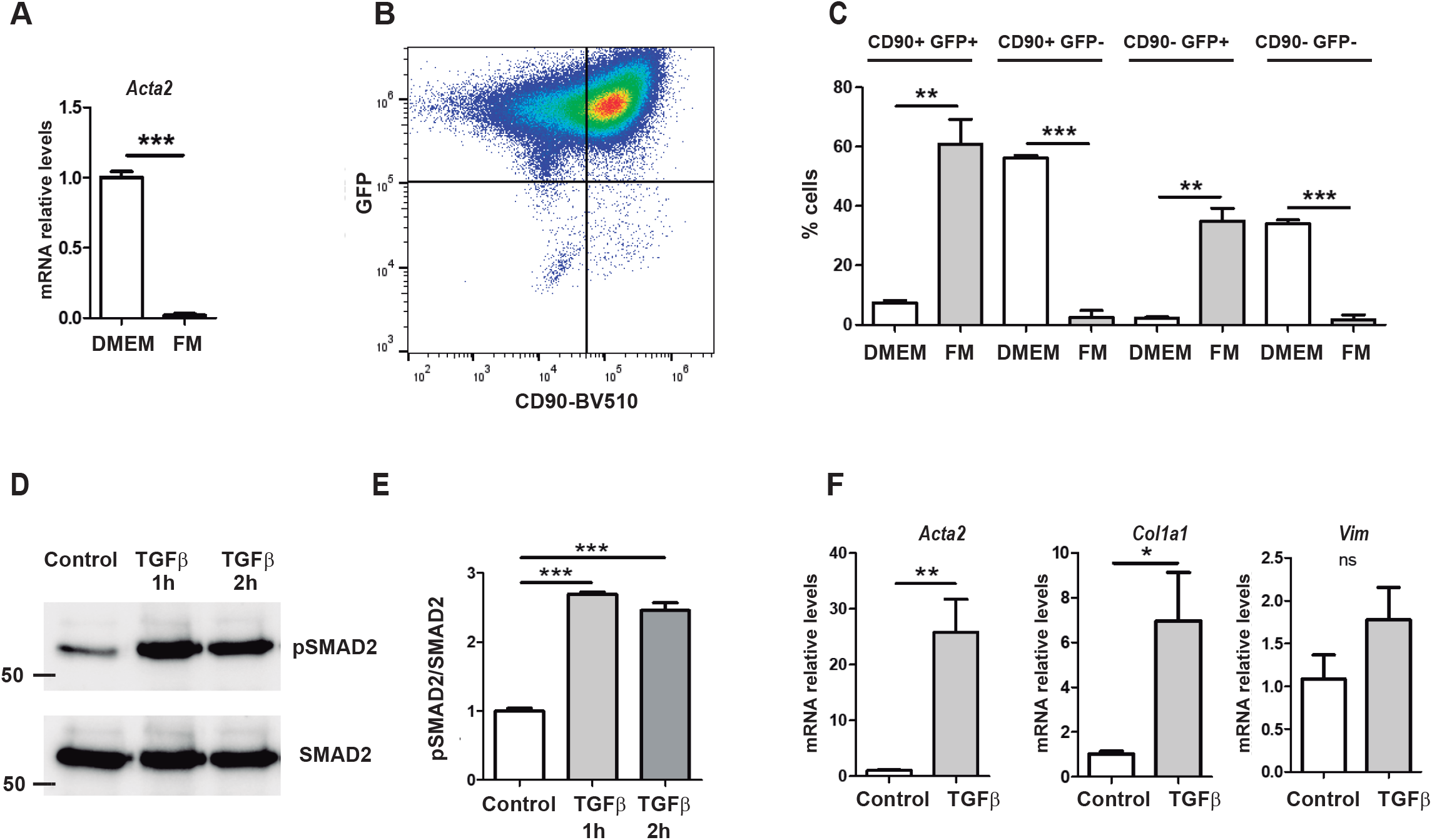
TGF-β induces a profibrotic phenotype in RFP^+^ cardiac fibroblasts. **(A)** qRT–PCR analysis of *Acta2* expression in RFP^+^ cardiac fibroblasts maintained in DMEM or fibroblast medium (FM). Data are presented as mean ± s.e.m. (n = 4); ***P < 0.0001 two-tailed student t-test. **(B)** Representative flow-cytometry plots of RFP^+^ cardiac fibroblasts cultured in FM. **(C)** Quantification of CD90/GFP fibroblast subsets (CD90^+^GFP^+^, CD90^+^GFP^−^, CD90^−^GFP^+^, CD90^−^GFP^−^) in DMEM versus FM. Data are presented as mean ± s.e.m. (n = 3), **P < 0.001, ***P < 0.0001, two-tailed student t-test (**D, E**) Western blot and densitometric analysis of SMAD2 phosphorylation (pSMAD2) in cardiac fibroblasts treated with TGF-β, with total SMAD2 serving as the loading control. Data are presented as mean ± s.e.m. (n = 3); ***P < 0.0001, one-way ANOVA followed by Tukey’s post-hoc test. (**F)** qRT– PCR analysis of the indicated genes in cardiac fibroblasts cultured with TGF-β for 3 days. Data are presented as mean ± s.e.m. (n = 3), *P < 0.05, **P < 0.001, two-tailed student t-test.

After successfully controlling the activation of our cell culture, our next objective was to investigate the effect of TGF-β, a central regulator of tissue fibrosis, and assess whether our model could serve as an ideal system for studying cardiac fibrosis in vitro (Deng et al, 2024). Western blot analysis revealed a robust increase in Smad2 phosphorylation in response to TGF-β, confirming the activation of the TGF-β-SMAD signaling cascade (Fig. 4D, E). To further characterize the fibrotic response of CFs to TGF-β, we examined the expression of two key fibrosis markers, *Acta2* and *Col1a1*. qRT-PCR analysis demonstrated a significant upregulation of *Acta2* and *Col1a1* expression in TGF-β-treated CFs, highlighting the induction of a fibrotic phenotype. In contrast, the expression of *Vim*, a pan-fibroblast marker, remained unchanged, suggesting that the observed changes were specific to a fibrotic phenotype (Fig. 4F).

These findings collectively demonstrate that the immortalized CFs constitute a robust and reliable model for studying stimulus-induced fibrotic activation in CFs.

## DISCUSSION

We developed a novel mouse model that enables distinct cardiac stromal cell populations, including resident fibroblasts, to be isolated based on WT1 reporter expression. The Wt1^GFP/+^;Wt1Cre;ROSA26-tdRFP model facilitates both the lineage tracing of cells labeled with Wt1Cre (RFP^+^) and the identification of cells actively expressing WT1 (GFP^+^). We generated an immortalized population highly enriched in different CF subtypes that retained RFP expression and expressed key fibroblast-specific markers, such as vimentin. The integrated GFP reporter enabled the isolation and functional analysis of distinct subpopulations within the Wt1Cre lineage.

During cardiac development, a subset of epicardial cells undergoes epithelial-to-mesenchymal transition, giving rise to EPDCs. These cells invade the subepicardial matrix before migrating into the myocardium, where they differentiate into various cardiac lineages, including most CFs (Gittenberger-de Groot et al, 1998; von Gise & Pu, 2012). During early postnatal (P) development, CFs undergo significant expansion via cell proliferation, which dramatically decreases after P7 (Ieda et al, 2009; Wu et al, 2020). While several mechanisms that regulate the initial formation and differentiation of EPDCs have been elucidated, the molecular pathways controlling CF proliferation and terminal differentiation remain largely unknown (Tallquist, 2020; von Gise & Pu, 2012). Given these limitations, the immortalized CFs generated in this study offer a valuable platform for screening candidate pathways involved in fibroblast function. Indeed, such analyses are typically difficult to perform using primary cultures because they contain limited numbers of cells in each identified subpopulation, hindering in-depth cellular and molecular studies.

The utility of this model also extends beyond developmental studies, because the adult mammalian heart has a limited capacity for regeneration. Healing typically occurs through scar formation, in which resident fibroblasts of epicardial origin play a central role (Tallquist, 2020). Although this response is essential for maintaining structural integrity and preventing ventricular rupture following injury, excessive or dysregulated activation can result in adverse cardiac remodeling, reduced contractile function, and ultimately, heart failure (Moore-Morris & Evans, 2023; Quijada et al, 2019). Notably, gene expression profiling has revealed that proliferating fibroblasts following MI closely resemble neonatal fibroblasts underscoring the relevance of our model (Kretzschmar et al, 2018). Understanding the regulatory mechanisms that govern the expansion and differentiation of CFs into specific subpopulations could help develop therapeutic strategies that mitigate the effects of pathological fibrosis while preserving essential wound healing processes.

WT1 is a transcription factor essential for development and tissue homeostasis (Hastie, 2017). Recent studies have highlighted its profibrotic role in lung fibrosis; however, WT1 deletion in the liver has also been shown to exacerbate fibrogenesis following injury (Kendall et al, 2019; Sontake et al, 2018). In the heart, studies using a zebrafish model of myocardial injury have shown that *wt1a:GFP^+^* cells upregulate a fibrotic gene program in response to damage. After analyzing different scRNA-seq datasets in mammals, WT1 has been classified as an early transcriptional regulator following myocardial infarction (Patrick et al, 2024; Sanchez-Iranzo et al, 2018). Despite these findings, the contribution of Wt1-expressing fibroblasts to the early stages of fibroblast activation and scar tissue formation after myocardial infarction remains unclear. Interestingly, CD90^+^GFP^−^ cells exhibited higher *Acta2* expression than CD90^+^GFP^+^ cells. Based on this, we hypothesize that, similar to TCF21, WT1 functions as a repressor of fibroblast activation and acquisition of a myofibroblast phenotype (Johansen et al, 2025).

Currently, no effective pharmacological treatment for cardiac fibrosis exists, posing a major challenge in cardiovascular medicine. Our immortalized cells represent the first specific model of epicardial-origin CFs, offering a unique and valuable platform for studying the molecular mechanisms underlying cardiac fibrosis. Given their robustness and relevance, our model could prove highly effective for high-throughput drug screening, aiding in the identification of novel therapeutic agents that may reverse or alleviate cardiac fibrosis.

Furthermore, the ability to genetically modify these cells facilitates the development of reporter-specific models, providing deeper insights into the cellular and molecular processes driving fibrosis in the heart and paving the way for more targeted, effective treatments.

In summary, we have developed a versatile model for studying CFs of epicardial origin. By adjusting the culture conditions, specific fibroblast subpopulations can be selectively promoted to enhance the model’s flexibility. The choice of conditions should be guided by the physiologically relevant context and desired subpopulation, which will allow for not only screening GFP expression but also for identifying treatment combinations that can modulate the proportions of different cell subpopulations. In turn, this can provide deeper insights into the functional roles of these cells.

## MATERIAL AND METHODS

### Animal models

The Wt1^GFP/+^, Wt1Cre (Tg(Wt1-cre)#Jbeb), and ROSA26-tdRFP mice have been described previously (Hosen et al, 2007; Luche et al, 2007; Wessels et al, 2012). To generate Wt1Cre;ROSA26-tdRFP mice, male Wt1Cre mice were mated with ROSA26dtRFP females. To generate Wt1^GFP/+^;Wt1Cre;ROSA26-tdRFP mice, Wt1Cre;ROSA26-tdRFP males were mated with Wt1^GFP/+^ females. The primers used for genotyping are listed in Table 7.

**Table 7.**
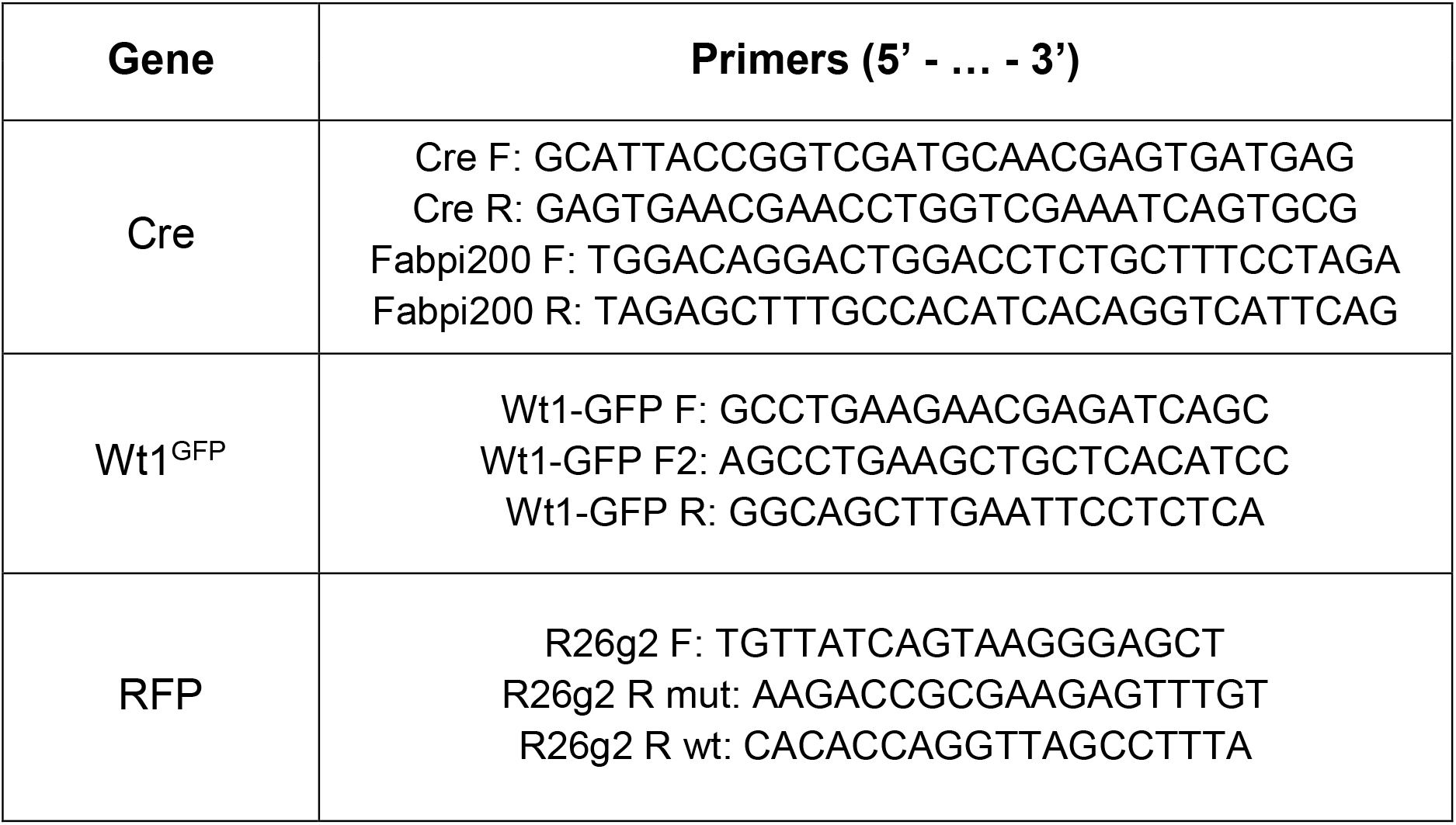
List of PCR primers used for genotyping

All animal experiments were carried out in accordance with the regulations of the Animal Experimentation Ethics Committee (CEEA) of the University of Barcelona (ID C121CG2V6), thereby complying with current Spanish and European legislation.

### Flow cytometry analysis

Flow cytometry analyses were performed on freshly isolated cells from the cardiac ventricles of mice or on immortalized cells.

For the in vivo analyses, early postnatal heart ventricles from Wt1^GFP/+^;Wt1Cre;ROSA26-tdRFP mice and littermate negative controls were digested in 1 mg/mL collagenase type I solution (Worthington, 9001-12-1) for 30 minutes at 37°C, using a shaking block set to 1000 rpm (Eppendorf Thermomixer Compact). Collagenase activity was stopped by washing the cells with Dulbecco’s Modified Eagle Medium (DMEM) (Gibco,11960044) containing 10% fetal bovine serum (FBS; Capricorn scientific, FBS-12B). Subsequently, the cells were then pelleted by centrifugation at 300*×g* for 5 minutes and filtered through a 40 μm cell strainer (Corning, 352340) before being resuspended in Hanks’ Balanced Salt Solution (HBSS Ca^2+^, Mg^2+^; Gibco, 14025092) containing 2% FBS (FACs buffer) (Ramiro-Pareta et al, 2023; Torres-Cano et al, 2022).

For antibody staining, freshly isolated or immortalized cells were incubated with primary monoclonal or isotype control antibodies at 4°C for 30 minutes (Table 8) (Ramiro-Pareta et al, 2023). Samples were washed twice and resuspended in FACs buffer. Flow cytometry analysis was carried out using a Cytek® Aurora 4L (Cytek) and data were analyzed using FlowJo™ software (v10.10). Cell gating of the corresponding fluorescent protein was carried out using samples negative for the fluorescent transgenic protein and isotype control antibodies.

**Table 8.**
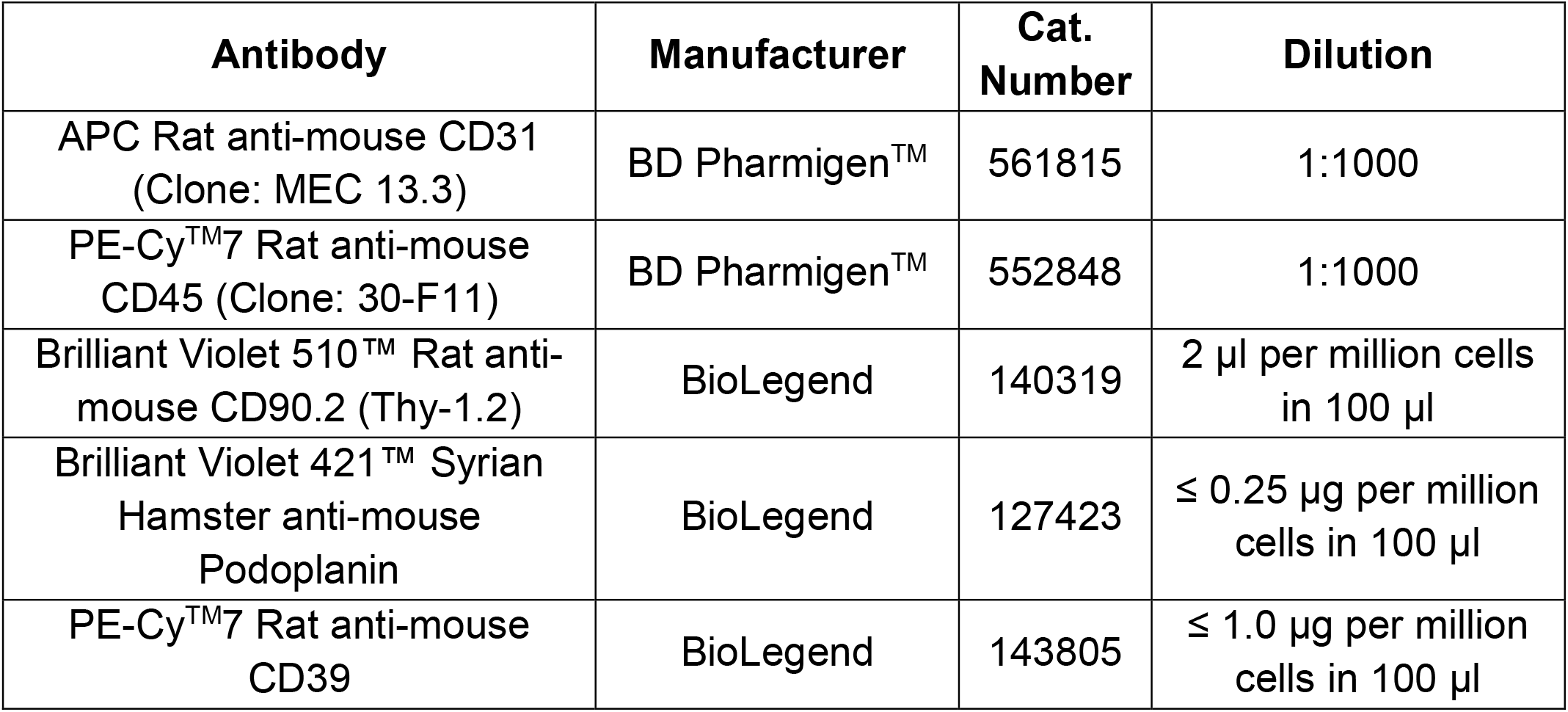
List of antibodies used for flow cytometry analysis

| Antibody | Manufacturer | Cat. Number | Dilution |
| --- | --- | --- | --- |
| APC Rat anti-mouse CD31<br>(Clone: MEC 13.3) | BD Pharmigen™ | 561815 | 1:1000 |
| PE-Cy™7 Rat anti-mouse<br>CD45 (Clone: 30-F11) | BD Pharmigen™ | 552848 | 1:1000 |
| Brilliant Violet 510™ Rat anti-mouse CD90.2 (Thy-1.2) | BioLegend | 140319 | 2 µl per million cells<br>in 100 µl |
| Brilliant Violet 421™ Syrian<br>Hamster anti-mouse<br>Podoplanin | BioLegend | 127423 | ≤ 0.25 µg per million<br>cells in 100 µl |
| PE-Cy™7 Rat anti-mouse<br>CD39 | BioLegend | 143805 | ≤ 1.0 µg per million<br>cells in 100 µl |

### Immortalization of RFP+ cardiac cells

RFP^+^ cells were isolated by FACS (FACSAria™ Fusion I, BD) from the enzymatically digested hearts of Wt1^GFP/+^;Wt1Cre;ROSA26-tdRFP mice, excluding endothelial (RFP^+^, CD31^+^, CD45^−^), hematopoietic (RFP^+^, CD31^−^, CD45^+^), and epicardial (RFP^+^, GFP^++^) cell populations. The sorted RFP^+^ cells were subsequently cultured on 24-well plates coated with poly-L-lysine (Sigma, P7886) in completed cell culture medium (DMEM high-glucose; Gibco, 11960044) supplemented with L-glutamine (Corning, 25-005-CV), 100 µg/mL streptomycin, 100 U/mL penicillin (Gibco, 15140122), and 10% FBS (Capricorn Scientific, FBS-12B) until reaching approximately 70% confluence.

To induce immortalization, cells were incubated overnight at 37°C with 1.2 µL of SV40 viral vector supernatant (GeneCopoeia, LP721-025). After 24 hours, the supernatant was removed and fresh culture medium was added. Cells were then cultured for an additional 21 days in presence of puromycin (Sigma, P8833). Non-transduced cells served as controls for the immortalization process. Once confluence was achieved, cells were trypsinized and designated as passage one (p1).

### Fibroblast medium and TGF-β treatment conditions

Immortalized cells were maintained in complete DMEM culture medium and passaged every 4 to 5 days upon reaching 70% confluency. To assess the impact of fibroblast medium on cellular phenotype and marker expression stability, cells were cultured in a commercially available fibroblast medium (Promocell, C-23025) for three passages, followed by subsequent analyses. Cells between passages 6 and 16 were used for all experiments.

To investigate the effect of TGF-β stimulation, cells were cultured at a density of 10,000 cells/cm^2^ in fibroblast growth medium for 24 hours, followed by overnight incubation in Opti-MEM medium. TGF-β (10 ng/mL) was then applied for 1, 2 or 72 hours, after which samples were collected for marker analysis by Western blot or qPCR, respectively.

### Isolation of RNA and real-time PCR

Total RNA from mixed cultures or FACS-sorted subpopulations was isolated using the PureLink RNA Mini Kit (Invitrogen, Cat. No. 12183025) according to the manufacturer’s instructions. The purified RNA was reverse transcribed into cDNA using UltraScript® Reverse Transcriptase (PCR Biosystems, PB30.11-10). Relative gene expression levels were quantified using SYBR Master Mix (Promega, A6002) and real-time PCR was performed on 96-well QuantStudio 3 (AppliedBiosystems) plates using the Connect analysis software (Thermo Fisher). The primers used are listed in Table 9.

**Table 9.**
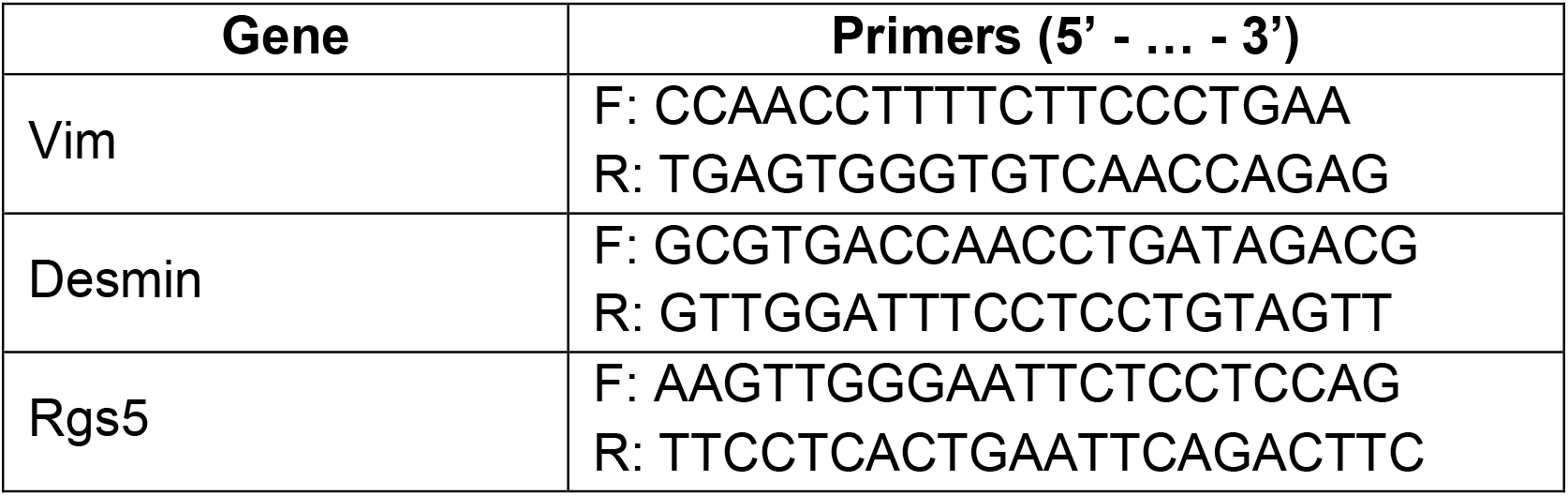
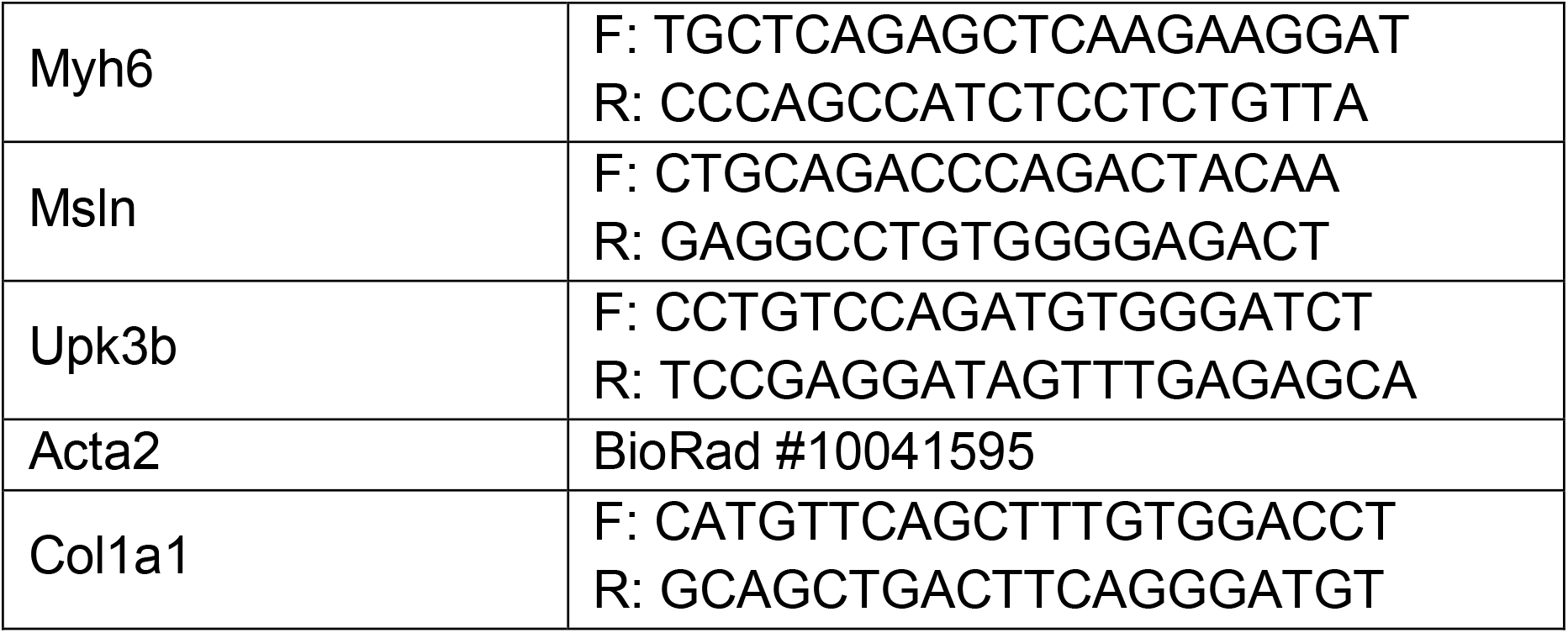
qRT-PCR primer list

### Immunofluorescence

Immortalized cells were seeded on 12 mm glass coverslips (6,000 cells/cm^2^) and cultured for 48 hours in complete DMEM culture medium or fibroblast growth medium. Cells were then fixed with 4% paraformaldehyde (Sigma, 158127) for 15 minutes, washed with phosphate buffered saline (PBS), permeabilized with 0.1% Triton X-100 (Sigma, 9036-19-5) for 5 minutes, washed again with PBS, and blocked with PBS supplemented with 1% bovine serum albumin (Sigma, 810664) for 1 hour at room temperature. After blocking, cells were incubated overnight at 4°C with primary antibodies against vimentin and SMA along with phalloidin, to stain filamentous actin (Table 10). Then, the cells were incubated with fluorochrome-conjugated secondary antibodies (Table 10) for 1 hour at 37°C. Cell nuclei were then stained with DAPI (Sigma, D9542). Immunofluorescence images were obtained using a fluorescence microscope (Leica Thunder).

**Table 10.** List of primary and secondary antibodies used for immunofluorescence.

| **Antibody** | **Manufacturer** | **Cat. Number** | **Species** | **Dilution** |
| --- | --- | --- | --- | --- |
| Anti-Vimentin | Abcam | ab8978 | Mouse | 1:100 |
| Anti-αSMA | Cell Signalling | 19245S | Rabbit | 1:100 |
| Phalloidin-TRITC | Sigma | P1951 |  | 1:200 |
| Anti-mouse, AlexaFluor 546 | Invitrogen | A11003 | Goat | 1:400 |
| Anti-rabbit, Alexa Fluor 633 | Invitrogen | A21070 | Goat | 1:400 |

### Western Blot

For western blot analyses, control and cells treated with TGF-β (10 ng/mL for 1 and 2 hours) were lysed using SDS buffer (1.5 M Tris pH 6.8, 15% glycerol, 3% SDS, 7.5% β-mercaptoethanol and 0.0375% Bromophenol Blue). Lysates were separated by SDS-PAGE gel and transferred to PVDF membranes for immunoblotting. Membranes were blocked for 1 h at room temperature in Tris-buffered saline containing 0.1% Tween-20 (TBST) and 5% bovine serum albumin (BSA) and immunoblots were incubated overnight at 4°C with antibodies against p-Smad2 and Smad2. Signal detection was performed using HRP-conjugated IgG secondary antibodies (Table 11). Blot intensities were quantified using ImageJ software and one-way ANOVA followed by Tukey’s post hoc test was used to examine statistical significance.

**Table 11.** List of antibodies used for Western Blot analysis.

| **Antibody** | **Manufacturer** | **Cat. Number** | **Species** | **Dilution** |
| --- | --- | --- | --- | --- |
| p-SMAD2 | Cell Signalling | 3108 | Rabbit | 1:1000 |
| SMAD2 | Cell Signalling | 3103 | Mouse | 1:1000 |
| Anti-Rabbit IgG HRP conjugate | BioRad | 1706515 | Goat | 1:3000 |
| Anti-Mouse IgG HRP conjugate | BioRad | 1706516 | Goat | 1:3000 |

### Quantification and statistical analysis

Data are presented as mean ± standard error of the mean (s.e.m). Statistical significance between two groups was determined using an unpaired, two-tailed Student’s t-test. For comparisons among multiple groups, one-way ANOVA followed by Tukey’s post hoc test was applied. Statistical analyses were performed using GraphPad Prism version 5 (GraphPad Software).

## Acknowledgments

We are grateful for the assistance provided by the Centres Científics i Tecnològics (CCiTUB) of the University of Barcelona for the use of advanced microscopy, flow cytometry, and animal facilities.

## References

Acharya A, Baek ST, Banfi S, Eskiocak B, Tallquist MD (2011) Efficient inducible Cre-mediated recombination in Tcf21 cell lineages in the heart and kidney. Genesis 49: 870–877

Acharya A, Baek ST, Huang G, Eskiocak B, Goetsch S, Sung CY, Banfi S, Sauer MF, Olsen GS, Duffield JS, Olson EN, Tallquist MD (2012) The bHLH transcription factor Tcf21 is required for lineage-specific EMT of cardiac fibroblast progenitors. Development 139: 2139–2149

Braitsch CM, Kanisicak O, van Berlo JH, Molkentin JD, Yutzey KE (2013) Differential expression of embryonic epicardial progenitor markers and localization of cardiac fibrosis in adult ischemic injury and hypertensive heart disease. Journal of molecular and cellular cardiology 65: 108–119

Cano E, Carmona R, Ruiz-Villalba A, Rojas A, Chau YY, Wagner KD, Wagner N, Hastie ND, Munoz-Chapuli R, Perez-Pomares JM (2016) Extracardiac septum transversum/proepicardial endothelial cells pattern embryonic coronary arterio-venous connections. Proceedings of the National Academy of Sciences of the United States of America 113: 656–661

Deng Y, He Y, Xu J, He H, Zhang M, Li G (2025) Cardiac fibroblasts regulate myocardium and coronary vasculature development in the murine heart via the collagen signaling pathway. eLife 13

Deng Z, Fan T, Xiao C, Tian H, Zheng Y, Li C, He J (2024) TGF-beta signaling in health, disease, and therapeutics. Signal transduction and targeted therapy 9: 61

Feng W, Bais A, He H, Rios C, Jiang S, Xu J, Chang C, Kostka D, Li G (2022) Single-cell transcriptomic analysis identifies murine heart molecular features at embryonic and neonatal stages. Nature communications 13: 7960

Forte E, Skelly DA, Chen M, Daigle S, Morelli KA, Hon O, Philip VM, Costa MW, Rosenthal NA, Furtado MB (2020) Dynamic Interstitial Cell Response during Myocardial Infarction Predicts Resilience to Rupture in Genetically Diverse Mice. Cell reports 30: 3149–3163 e3146

Garvin AM, Katwa LC (2023) Primary cardiac fibroblast cell culture: methodological considerations for physiologically relevant conditions. American journal of physiology Heart and circulatory physiology 325: H869–H881

Gittenberger-de Groot AC, Vrancken Peeters MP, Mentink MM, Gourdie RG, Poelmann RE (1998) Epicardium-derived cells contribute a novel population to the myocardial wall and the atrioventricular cushions. Circulation research 82: 1043–1052

Hamilton TG, Klinghoffer RA, Corrin PD, Soriano P (2003) Evolutionary divergence of platelet-derived growth factor alpha receptor signaling mechanisms. Molecular and cellular biology 23: 4013–4025

Hastie ND (2017) Wilms’ tumour 1 (WT1) in development, homeostasis and disease. Development 144: 2862–2872

Hortells L, Valiente-Alandi I, Thomas ZM, Agnew EJ, Schnell DJ, York AJ, Vagnozzi RJ, Meyer EC, Molkentin JD, Yutzey KE (2020) A specialized population of Periostin-expressing cardiac fibroblasts contributes to postnatal cardiomyocyte maturation and innervation. Proceedings of the National Academy of Sciences of the United States of America 117: 21469–21479

Hosen N, Shirakata T, Nishida S, Yanagihara M, Tsuboi A, Kawakami M, Oji Y, Oka Y, Okabe M, Tan B, Sugiyama H, Weissman IL (2007) The Wilms’ tumor gene WT1-GFP knock-in mouse reveals the dynamic regulation of WT1 expression in normal and leukemic hematopoiesis. Leukemia 21: 1783–1791

Humeres C, Frangogiannis NG (2019) Fibroblasts in the Infarcted, Remodeling, and Failing Heart. JACC Basic to translational science 4: 449–467

Ieda M, Tsuchihashi T, Ivey KN, Ross RS, Hong TT, Shaw RM, Srivastava D (2009) Cardiac fibroblasts regulate myocardial proliferation through beta1 integrin signaling. Developmental cell 16: 233–244

Johansen AKZ, Kasam RK, Vagnozzi RJ, Lin SJ, Gomez-Arroyo J, Shittu A, Bowers SLK, Kuwabara Y, Grimes KM, Warrick K, Sargent MA, Baldwin TA, Quaggin SE, Barski A, Molkentin JD (2025) Transcription Factor 21 Regulates Cardiac Myofibroblast Formation and Fibrosis. Circulation research 136: 44–58

Kendall TJ, Duff CM, Boulter L, Wilson DH, Freyer E, Aitken S, Forbes SJ, Iredale JP, Hastie ND (2019) Embryonic mesothelial-derived hepatic lineage of quiescent and heterogenous scar-orchestrating cells defined but suppressed by WT1. Nature communications 10: 4688

Kretzschmar K, Post Y, Bannier-Helaouet M, Mattiotti A, Drost J, Basak O, Li VSW, van den Born M, Gunst QD, Versteeg D, Kooijman L, van der Elst S, van Es JH, van Rooij E, van den Hoff MJB, Clevers H (2018) Profiling proliferative cells and their progeny in damaged murine hearts. Proceedings of the National Academy of Sciences of the United States of America 115: E12245–E12254

Luche H, Weber O, Nageswara Rao T, Blum C, Fehling HJ (2007) Faithful activation of an extra-bright red fluorescent protein in “knock-in” Cre-reporter mice ideally suited for lineage tracing studies. European journal of immunology 37: 43–53

Martinez-Estrada OM, Lettice LA, Essafi A, Guadix JA, Slight J, Velecela V, Hall E, Reichmann J, Devenney PS, Hohenstein P, Hosen N, Hill RE, Munoz-Chapuli R, Hastie ND (2010) Wt1 is required for cardiovascular progenitor cell formation through transcriptional control of Snail and E-cadherin. Nature genetics 42: 89–93

Moore-Morris T, Cattaneo P, Guimaraes-Camboa N, Bogomolovas J, Cedenilla M, Banerjee I, Ricote M, Kisseleva T, Zhang L, Gu Y, Dalton ND, Peterson KL, Chen J, Puceat M, Evans SM (2018) Infarct Fibroblasts Do Not Derive From Bone Marrow Lineages. Circulation research 122: 583–590

Moore-Morris T, Evans SM (2023) Cardiac Pericyte Diversity in Infarct Remodeling: Not Just Vascular Support Cells? Circulation 148: 899–902

Moore-Morris T, Guimaraes-Camboa N, Banerjee I, Zambon AC, Kisseleva T, Velayoudon A, Stallcup WB, Gu Y, Dalton ND, Cedenilla M, Gomez-Amaro R, Zhou B, Brenner DA, Peterson KL, Chen J, Evans SM (2014) Resident fibroblast lineages mediate pressure overload-induced cardiac fibrosis. The Journal of clinical investigation 124: 2921–2934

Moore AW, McInnes L, Kreidberg J, Hastie ND, Schedl A (1999) YAC complementation shows a requirement for Wt1 in the development of epicardium, adrenal gland and throughout nephrogenesis. Development 126: 1845–1857

Patrick R, Janbandhu V, Tallapragada V, Tan SSM, McKinna EE, Contreras O, Ghazanfar S, Humphreys DT, Murray NJ, Tran YTH, Hume RD, Chong JJH, Harvey RP (2024) Integration mapping of cardiac fibroblast single-cell transcriptomes elucidates cellular principles of fibrosis in diverse pathologies. Science advances 10: eadk8501

Perez-Pomares JM, Phelps A, Sedmerova M, Carmona R, Gonzalez-Iriarte M, Munoz-Chapuli R, Wessels A (2002) Experimental studies on the spatiotemporal expression of WT1 and RALDH2 in the embryonic avian heart: a model for the regulation of myocardial and valvuloseptal development by epicardially derived cells (EPDCs). Developmental biology 247: 307–326

Pogontke C, Guadix JA, Sanchez-Tevar AM, Munoz-Chapuli R, Ruiz-Villalba A, Perez-Pomares JM (2022) Dynamic Epicardial Contribution to Cardiac Interstitial c-Kit and Sca1 Cellular Fractions. Frontiers in cell and developmental biology 10: 864765

Quijada P, Misra A, Velasquez LS, Burke RM, Lighthouse JK, Mickelsen DM, Dirkx RA, Jr., Small EM (2019) Pre-existing fibroblasts of epicardial origin are the primary source of pathological fibrosis in cardiac ischemia and aging. Journal of molecular and cellular cardiology 129: 92–104

Ramiro-Pareta M, Muller-Sanchez C, Portella-Fortuny R, Soler-Botija C, Torres-Cano A, Esteve-Codina A, Bayes-Genis A, Reina M, Soriano FX, Montanez E, Martinez-Estrada OM (2023) Endothelial deletion of Wt1 disrupts coronary angiogenesis and myocardium development. Development 150

Ruiz-Villalba A, Simon AM, Pogontke C, Castillo MI, Abizanda G, Pelacho B, Sanchez-Dominguez R, Segovia JC, Prosper F, Perez-Pomares JM (2015) Interacting resident epicardium-derived fibroblasts and recruited bone marrow cells form myocardial infarction scar. Journal of the American College of Cardiology 65: 2057–2066

Sanchez-Iranzo H, Galardi-Castilla M, Sanz-Morejon A, Gonzalez-Rosa JM, Costa R, Ernst A, Sainz de Aja J, Langa X, Mercader N (2018) Transient fibrosis resolves via fibroblast inactivation in the regenerating zebrafish heart. Proceedings of the National Academy of Sciences of the United States of America 115: 4188–4193

Sontake V, Kasam RK, Sinner D, Korfhagen TR, Reddy GB, White ES, Jegga AG, Madala SK (2018) Wilms’ tumor 1 drives fibroproliferation and myofibroblast transformation in severe fibrotic lung disease. JCI insight 3

Tallquist MD (2018) Cardiac fibroblasts: from origin to injury. Current opinion in physiology 1: 75–79

Tallquist MD (2020) Cardiac Fibroblast Diversity. Annual review of physiology 82: 63–78

Tallquist MD, Molkentin JD (2017) Redefining the identity of cardiac fibroblasts. Nature reviews Cardiology 14: 484–491

Torres-Cano A, Portella-Fortuny R, Muller-Sanchez C, Porras-Marfil S, Ramiro-Pareta M, Chau YY, Reina M, Soriano FX, Martinez-Estrada OM (2022) Deletion of Wt1 during early gonadogenesis leads to differences of sex development in male and female adult mice. PLoS genetics 18: e1010240

Velecela V, Torres-Cano A, Garcia-Melero A, Ramiro-Pareta M, Muller-Sanchez C, Segarra-Mondejar M, Chau YY, Campos-Bonilla B, Reina M, Soriano FX, Hastie ND, Martinez FO, Martinez-Estrada OM (2019) Epicardial cell shape and maturation are regulated by Wt1 via transcriptional control of Bmp4. Development 146

von Gise A, Pu WT (2012) Endocardial and epicardial epithelial to mesenchymal transitions in heart development and disease. Circulation research 110: 1628–1645

Wessels A, van den Hoff MJ, Adamo RF, Phelps AL, Lockhart MM, Sauls K, Briggs LE, Norris RA, van Wijk B, Perez-Pomares JM, Dettman RW, Burch JB (2012) Epicardially derived fibroblasts preferentially contribute to the parietal leaflets of the atrioventricular valves in the murine heart. Developmental biology 366: 111–124

Wu R, Ma F, Tosevska A, Farrell C, Pellegrini M, Deb A (2020) Cardiac fibroblast proliferation rates and collagen expression mature early and are unaltered with advancing age. JCI insight 5

Yata Y, Scanga A, Gillan A, Yang L, Reif S, Breindl M, Brenner DA, Rippe RA (2003) DNase I-hypersensitive sites enhance alpha1(I) collagen gene expression in hepatic stellate cells. Hepatology 37: 267–276

Zhang H, Huang X, Liu K, Tang J, He L, Pu W, Liu Q, Li Y, Tian X, Wang Y, Zhang L, Yu Y, Wang H, Hu R, Wang F, Chen T, Wang QD, Qiao Z, Zhang L, Lui KO, Zhou B (2017) Fibroblasts in an endocardial fibroelastosis disease model mainly originate from mesenchymal derivatives of epicardium. Cell research 27: 1157–1177

Zhang LP, Nie Q, Kang JB, Wang B, Cai CL, Li JG, Qi WJ (2008) [Efficacy of whole body gamma-knife radiotherapy combined with thermochemotherapy on locally advanced pancreatic cancer]. Ai Zheng 27: 1204–1207

Zhou B, Honor LB, He H, Ma Q, Oh JH, Butterfield C, Lin RZ, Melero-Martin JM, Dolmatova E, Duffy HS, Gise A, Zhou P, Hu YW, Wang G, Zhang B, Wang L, Hall JL, Moses MA, McGowan FX, Pu WT (2011) Adult mouse epicardium modulates myocardial injury by secreting paracrine factors. The Journal of clinical investigation 121: 1894–1904

Zhou B, Ma Q, Rajagopal S, Wu SM, Domian I, Rivera-Feliciano J, Jiang D, von Gise A, Ikeda S, Chien KR, Pu WT (2008) Epicardial progenitors contribute to the cardiomyocyte lineage in the developing heart. Nature 454: 109–113

